# Overruled by nature: A plastic response to an ecological regime shift disconnects a gene and its trait

**DOI:** 10.1101/2022.10.27.514021

**Authors:** F. Besnier, Ø. Skaala, V. Wennevik, F. Ayllon, K.R. Utne, P.T. Fjeldheim, K. Andersen-Fjeldheim, S. Knutar, K.A. Glover

**Affiliations:** Institute of Marine Research, Bergen, Norway; Institute of Biology, University of Bergen, Norway

## Abstract

In Atlantic salmon, age at maturation is a life history trait ruled by a sex-specific trade-off between reproductive success and survival. Following an ecological regime shift in 2005, many North Atlantic salmon populations currently display smaller size at age and delayed age at maturation. However, whether this change reflects rapid evolution or plastic response is unknown. Some 1500 historical and contemporary salmon from river Etne (Western Norway) genotyped at 50k SNPs revealed three loci significantly associated with age at maturation. These included *vgll3* and *six6*, which collectively explained 36 to 50% of the age at maturation variation in the 1983-1984 period. Strikingly, the combined influence of these genes was nearly absent in all samples from 2013-2016, despite allelic frequencies at *vgll3* remaining unchanged. We conclude that the regime shift has led to the sudden bypassing of the influence of *vgll3* and *six6* on maturation through growth-driven plasticity.

## Introduction

Understanding the mechanisms by which organisms adapt to their environments is a central question in biology (Losos 2000; Andrew *et al*. 2013). However, beyond the academic curiosity that motivates biologists to investigate how species evolved and adapted until now, accelerating climate change and ongoing habitat destruction catalyses a sense of urgency when considering the fate of many species in the future. As unprecedentedly fast environmental changes have occurred for many taxa during the last half century (Chevin *et al*. 2010; Dullinger *et al*. 2012; Pörtner *et al*. 2022), the current mechanisms of adaptation of many species may not couple with the pace at which environmental change is now occurring, leading thus to population declines and extinctions. Consequently, the mechanisms of adaptation and their potential to remain effective in a context of rapidly changing environment are becoming increasingly important to understand (García de Leániz *et al*. 2007; Candolin & Heuschele 2008; Kwan *et al*. 2008; Bandillo *et al*. 2017).

In the context of environmental change, genes associated with sexual conflicts (Parker 1974) are of particular interest. Sexual conflicts are associated with sex-specific selection and therefore expected to promote genetic diversity in the underlying genes (Rowe *et al*. 2018), thus potentially helping maintain standing genetic variation on important fitness traits within the population. An illustration of this occurs in anadromous Atlantic salmon (*Salmo salar* L.) where a single gene (*vgll3*) is strongly associated with the age at maturation, a highly adaptative trait subject to sexual conflict (Ayllon *et al*. 2015; Barson *et al*. 2015). In many species, age at maturation represents a trade-off, as late-maturation gives larger body sizes and thus higher reproductive success, but at the increased risk of dying before reproduction (Fleming & Einum 2011). In Atlantic salmon, the optimum value for this trait differs between sexes, as males mature earlier and smaller whereas females benefit from later maturation and a larger body size resulting in more and larger eggs (Fleming & Einum 2011). This sexual conflict is in part resolved by the mediation of *vgll3* (Barson *et al*. 2015) through sex-specific dominance.

Studies of *vgll3* and age of maturation in Atlantic salmon have resulted in inconstant estimations regarding the degree to which this gene influences the trait. Several studies have demonstrated a very strong association in wild North European populations (Ayllon *et al*. 2015; Barson *et al*. 2015; Czorlich *et al*. 2018; Jensen *et al*. 2022), but in stark contrast, little or no association has been observed in wild populations in North America (Boulding *et al*. 2019; Mohamed *et al*. 2019). Furthermore, conflicting observations have also been reported in domesticated Norwegian farmed strains reared under aquaculture conditions (Ayllon *et al*. 2019; Sinclair-Waters *et al*. 2020). This begs the question, why does the influence of *vgll3* on age at maturation vary so greatly?

Salmonids have been exposed to a wide range of anthropological challenges including habitat modifications since the Industrial Revolution (Forseth *et al*. 2017). Furthermore, during the past two decades, most salmon populations in the North Atlantic have shifted towards later maturation (Otero *et al*. 2012; Vollset *et al*. 2022) and smaller size at age (Quinn *et al*. 2006; Bal *et al*. 2017; Todd *et al*. 2021; Vollset *et al*. 2022). Although these changes are thought to be caused by a regime-shift in oceanic conditions in 2005 (Vollset *et al*. 2022), whether they reflect rapid evolution or plastic responses is unknown. The same trends towards later maturation have also been documented in the salmon population inhabiting the river Etne on the west coast of Norway, where the proportion of fish maturing after one year at sea dropped from 63% in the period 1983-84 to 34% in 2018-19 (Harvey *et al*. 2022). The main objective of the present study was to investigate whether the observed changes in age at maturation were the result of phenotypic plasticity, or alternatively, evolution in the gene(s) influencing this trait. To address this, we genotyped historical (early 1980’s – pre regime shift) and contemporary (mid 2010’s post regime shift) samples with a 50k panel of genome-wide SNPs. This approach revealed the dissociation between age at maturation and two loci, *vgll3* and *six6*, that explained 36 to 50% of the variation in the 1983-1984 period, but only 7% in the most recent samples.

## Material and Methods

### Samples

This study is based on samples obtained from adult salmon captured in the river Etne, in western Norway, 60° N. This river is home to a salmon population of typically 1000-2500 adults returning from the sea annually (Harvey *et al*. 2017). A permanent trapping facility installed in the river has permitted sampling almost the entire adult spawning population since 2014, which facilitated access to both an extensive set of DNA samples, as well as to phenotypic and phenological data. For the present study, 797 wild adult salmon captured in the 1983-84 angling season were compared to 751 wild adult salmon captured in the upstream fish trap in the period 2013 to 2016 (Besnier *et al*. 2022). For the contemporary samples (2013-2016), an estimation of individual genetic admixture was computed as the proportion of domestic ancestry in each individuals genome (see Besnier *et al*. (2022) for details). In addition, a sample of 350 domesticated farmed salmon escapees that were removed from the river in the period 1989-2012 (15 per year), were genotyped with the same set of markers.

### Genotyping and sex determination

All samples were genotyped on a ThermoFisher Axiom 57K single nucleotide polymorphism (SNP) array (NOFSAL03, 55735 markers) developed by Nofima (Norwegian institute for applied research in food aquaculture and fisheries) in collaboration with private aquaculture companies Mowi and SalmoBreed (Besnier *et al*. 2022). SNPs were checked following the “Best Practice Workflow” on the Affymetrix axiom analysis software (available at: https://www.thermofisher.com/no/en/home/technical-resources/software-downloads.html). SNPs with call rates lower than 0.97 and samples with call rates lower than 0.85 were discarded, whereas markers classified as “PolyHighResolution” (High resolution in both homozygous and heterozygous clusters) were conserved for further data analysis.

Fish were sexed by examining variants of the sdY gene (Yano *et al*. 2012; Eisbrenner *et al*. 2014); *i.e*., males were identified based on the presence of exons 2 and 4. Samples were genotyped on an Applied Biosystems ABI 3730 Genetic Analyser, and genotypes were called using GeneMapper (Applied Biosystems, v. 4.0). The analysis of the sdY gene provides an accurate identification of sex, although, a very low percentage of fish identified as genetic males are phenotypic females due to carrying an inactive pseudo-copy of the sdY gene with both exon 2 and 4 (Ayllon *et al*. 2020).

### Genome scan for loci associated with age at maturation

Age at maturation was modeled as a binary trait consisting of early maturing fish, *i.e*., fish returning after one sea-winter (1SW), also known as grisling, *vs*. late maturing fish, *i.e*., fish returning after two or more winters at sea (2^+^SW). The probability of maturing early was then modeled in a generalized linear model with logit link function:

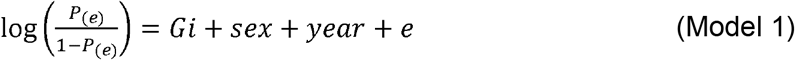

Where *P_(e)_* is the probability of early maturing, *G_i_* is the SNP genotype at locus *i, sex* is a binary factor accounting for genetically determined sex, *year* is a factor accounting for the sampling year, and *e* a normally distributed vector of residuals. Significance of association between genotype and sea age was estimated by comparing the deviance of Model 1 with the deviance of the model without genetic effect. The difference was compared to a Chi-squared distribution with one degree of freedom. Model 1 was fitted at each SNP available on the dataset, and the obtained p-values were adjusted for multiple testing by following Bonferroni correction. All p-values given in the genome scan result section are corrected for multiple testing.

### Haplotyping

Haplotypes were reconstructed using the Phase 2.1 software (Stephens *et al*. 2001) in the historical and contemporary data separately. Two loci on SSA9 and SSA25 were more specifically considered for haplotype reconstruction as they displayed high association with age at maturation, and associated genes *vgll3* and *six6* were previously described in the same two genomic regions (Ayllon *et al*. 2015; Barson *et al*. 2015; Czorlich *et al*. 2018). On SSA25, a haplotype window consisting of four polymorphic SNPs (AX-87309414, AX-172546510, AX-87309615, AX-87420691) was reconstructed in the region spanning from 28.65 to 28.66 Mb containing *vgll3* (https://www.ncbi.nlm.nih.gov/gene/106586514). On SSA09, a haplotype window consisting of three SNPs (AX-172546568, AX-88029383, AX-87668000), spanning from 24.86-24.95Mb was reconstructed around the position of *six6*. (https://www.ncbi.nlm.nih.gov/gene/106610974)

### Statistical analyses

#### Genetic structure to age at maturation variation

Due to the sex specific dominance observed in *vgll3*, the genetic structure was estimated for males and females separately. The probability of early maturation was modeled as a response to additive and dominance genetic effects as follow:

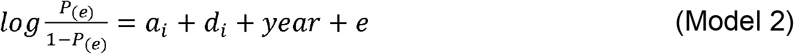

Where *P_(e)_* is the probability of early maturing, *a_i_* and *d_i_* are respectively the additive and dominance effects at locus *i*, *year* is a factor accounting for the sampling year, and *e* a normally distributed vector of residuals. The model was fitted in a GLM with logit link function in R (Team 2022), and the variance contribution was deduced from the difference in model deviance between Model 2 and a model than only accounted for capture year. Two loci interaction models were tested similarly by accounting for the additional interaction parameters between loci _i_ and _j_. In contrast with the genome scan, the test for significance if the different genetic parameters were not corrected for multiple testing. All p-values given with the estimation of genetic effect are nominal values.

#### Comparison in age at maturation

Potential differences in age at maturation between historical and contemporary samples was tested by chi-squared test on a contingency table reporting the number of observed 1, 2 and 3+ sea winter adults within historical and contemporary samples.

#### Size at age

Individual body length was recorded for every fish passing the trap in the contemporary sample, whereas the adult length of the 1983-1984 fish were calculated from reading scales samples. In addition, the growth at first sea winter was estimated for both historical and contemporary samples by calculating length at first sea winter from reading scales. The difference in length between samples from 1983-1984 and from 2013-2016 was tested with a two-sided t-test, separately for each sea-age and sex categories.

#### Potential role of admixture

The salmon population in the river Etne has been subject to introgression from domesticated salmon escaping from commercial fish farms, with an average 24% of genetic admixture in the contemporary population (Glover *et al*. 2013; Karlsson *et al*. 2016; Besnier *et al*. 2022). Individual admixture with domesticated salmon has already been correlated with earlier adult maturation in this population (Besnier *et al*. 2022). Therefore, in order to account for the potential influence of admixture on the temporal influence of loci on age at maturation in this population, we investigated whether the change in genetic architecture of age at maturation could be linked to genetic admixture.

A third model was fitted with the aim to evaluate a potential interaction between admixture and *vgll3* or *six6* genotypes.

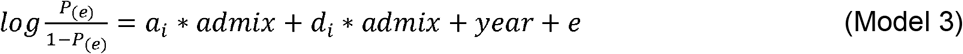

Where *admix* is the individual genetic admixture as computed in Besnier et al (Besnier *et al*. 2022).

## Results

### Temporal changes in age and size at maturation

Marine growth was compared between the historical (1983-84) and contemporary (2013-16) samples. During this period, the length of the fish during the first winter at sea decreased significantly, as well the length of the two sea-winter (2SW) adults, while the length of the one sea-winter (1SW) adults remained stable (Table 1).

**Table 1.** Comparison of fish length between historical and contemporary samples during first winter at sea (immature fish), adult maturing after one sea winter (1SW), and adult maturing after 2 winters at sea (2SW).

|  | phenotype | Mean value<br>(cm)<br>1983-1984 | Mean value<br>(cm)<br>2013-2016 | df | t | p |
| --- | --- | --- | --- | --- | --- | --- |
| Females | First sea<br>winter length | 47.2 | 41.7 | 500 | 14 | <0.001 |
|  | 1SW adult<br>length | 55.1 | 55.0 | 37 | 0.89 | 0.37 |
|  | 2SW adult<br>length | 79.4 | 75.3 | 202 | 5.50 | <0.001 |
| Males | First sea<br>winter length | 45.4 | 41.0 | 711 | 13.2 | <0.001 |
|  | 1SW adult<br>length | 55.6 | 56.8 | 441 | 2.7 | 0.007 |
|  | 2SW adult<br>length | 80.6 | 72.6 | 85 | 5.4 | <0.001 |

During the same period, we also observed a trend towards later maturation with a strong decline of the frequency of the fish maturing after one winter at sea from 46% to 8% for the females (χ^2^=120, df=1, p<2.10^−16^), and from 75% to 58% for the males (χ^2^=25, df=1, p=4.10^−7^) (Table 2).

**Table 2.** Percentage and numbers (in brackets) of fish displaying 1-2-3^+^ sea winter (SW) of age at maturation, by sex, in the historical (H – 1983/84, N = 797) and contemporary (C – 2013-2016, N = 751) samples.

|  | **1SW % (N)** | **2SW % (N)** | **3<sup>+</sup>SW % (N)** |
| --- | --- | --- | --- |
| Females_H | 46 <sub>(141)</sub> | 37 <sub>(114)</sub> | 17 <sub>(51)</sub> |
| Females_C | 8 <sub>(30)</sub> | 63 <sub>(227)</sub> | 28 <sub>(102)</sub> |
| Males_H | 75 <sub>(311)</sub> | 13 <sub>(56)</sub> | 12 <sub>(48)</sub> |
| Males_C | 58 <sub>(227)</sub> | 33 <sub>(127)</sub> | 9 <sub>(36)</sub> |

### Genome scans

The genome scan for age at maturation (Fig.1) identified three loci displaying a significant association in the historical samples, one on each of chromosomes SSA09, SSA24 and SSA25. The genomic regions identified in SSA25 and SSA09 overlapped with loci previously described as major contributors to the sea age variability in salmon: *vgll3* on SSA25 and *six6* located on SSA09. The degree of association between age at maturation and these genomic regions was investigated further in the historical and contemporary dataset.

**Figure 1:**
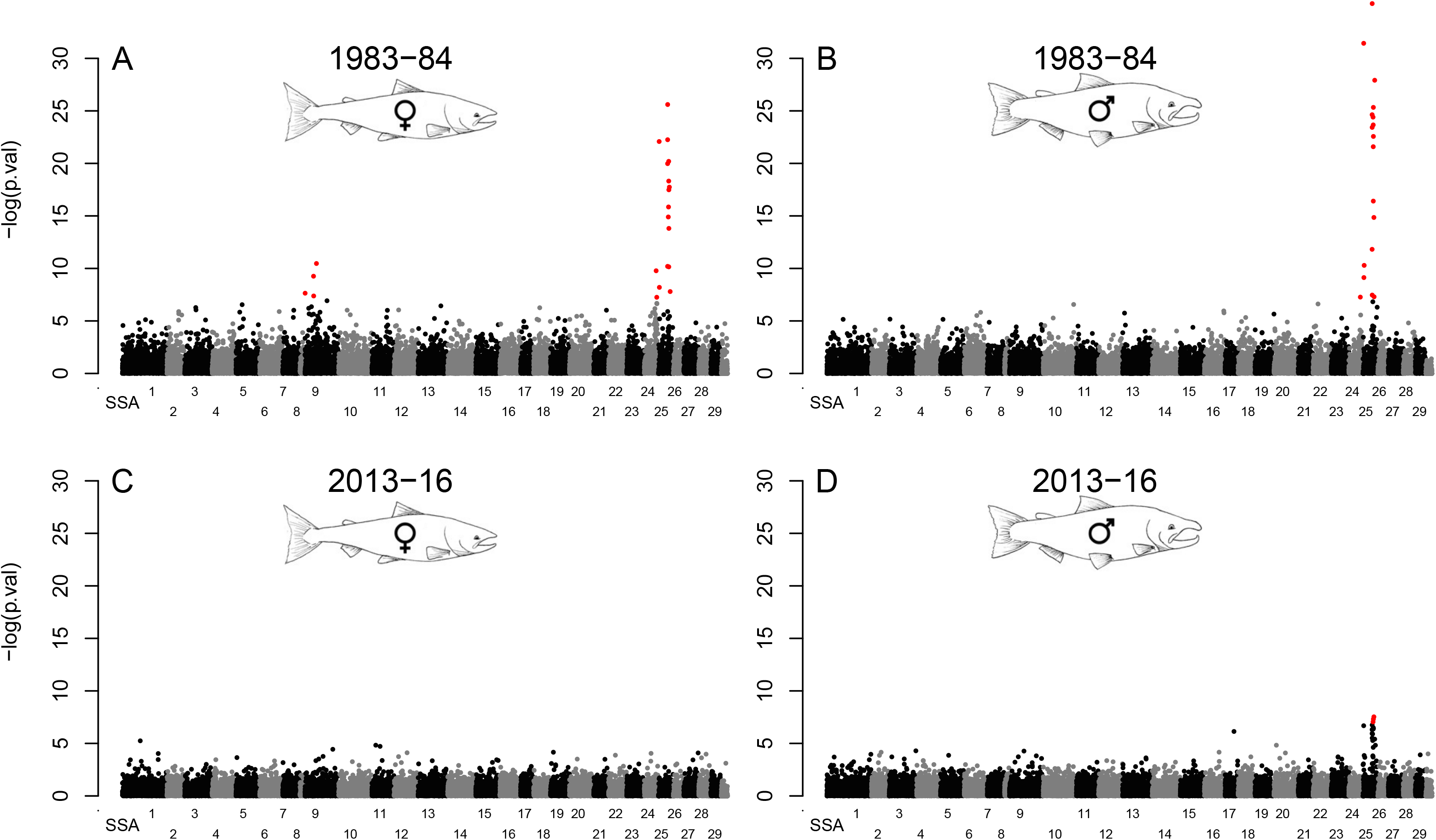
Scan for SNP association with age at maturation. Red dots represent SNPs with significant association with sea age (p<0.01 after correction for multiple tests). Females from 1983-84 (**A**), males from 1983-84 (**B**), females from 2013-16 (**C**) and males from 2013-16 (**D**).

### *vgll3* and *six6* haplotypes

Haplotypes were reconstructed in the historical and contemporary samples separately, across the genomic regions in SSA25 and SSA9 that contained the *vgll3* and s*ix6* genes respectively. On SSA25, two haplotypes were predominant in the historical samples with frequencies of 56% and 39% (Table 3) and respectively associated with early and late sea age. The mean sea age was 1.18 for the homozygous haplotype “1221”, and 2.46 for the homozygous haplotype “2112”, which will thus be referred to as *vgll3*-E and *vgll3*-L alleles from hereon. In the contemporary samples, the same two haplotypes were found in almost identical frequencies (df=1, χ^2^=0.85, p=0.35) to the historical samples (59% and 39% for *vgll3*-E and *vgll3*-L respectively), indicating that no temporal change in haplotype frequencies occurred at this locus during the three-decade period.

**Table 3.**
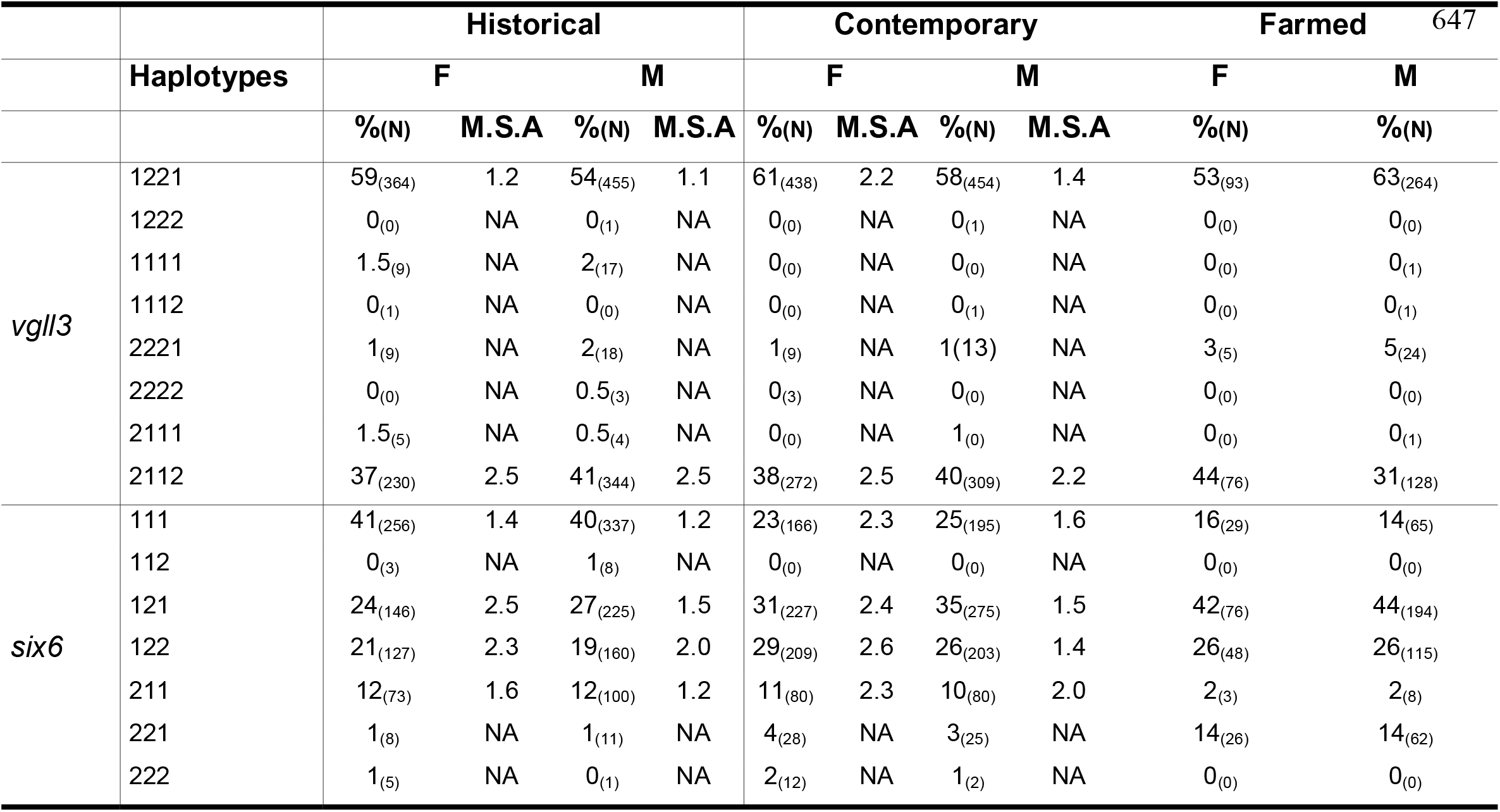
Observed occurrences of each haplotype per sex and associated Mean Sea Age (M.S.A) in the historical, contemporary and farmed samples. (M.S.A is calculated as mean sea age of the samples that are homozygous for the haplotype.)

The mean age at maturation in each genotype class (Fig. 2) displayed a strong association with an additive effect of *vgll3* on female sea age in the historical data (Fig. 2.A, Table.S1), whereas association of *vgll3* almost disappeared in the contemporary female data (Fig. 2.C, Table.S1). For the historical male data (Fig. 2B, Table.S1), a strong *vgll3* association with a dominant *vgll3*-E allele was also detected, whereas the genetic association was strongly reduced in the contemporary data (Fig. 2.D, Table.S1).

**Figure 2:**
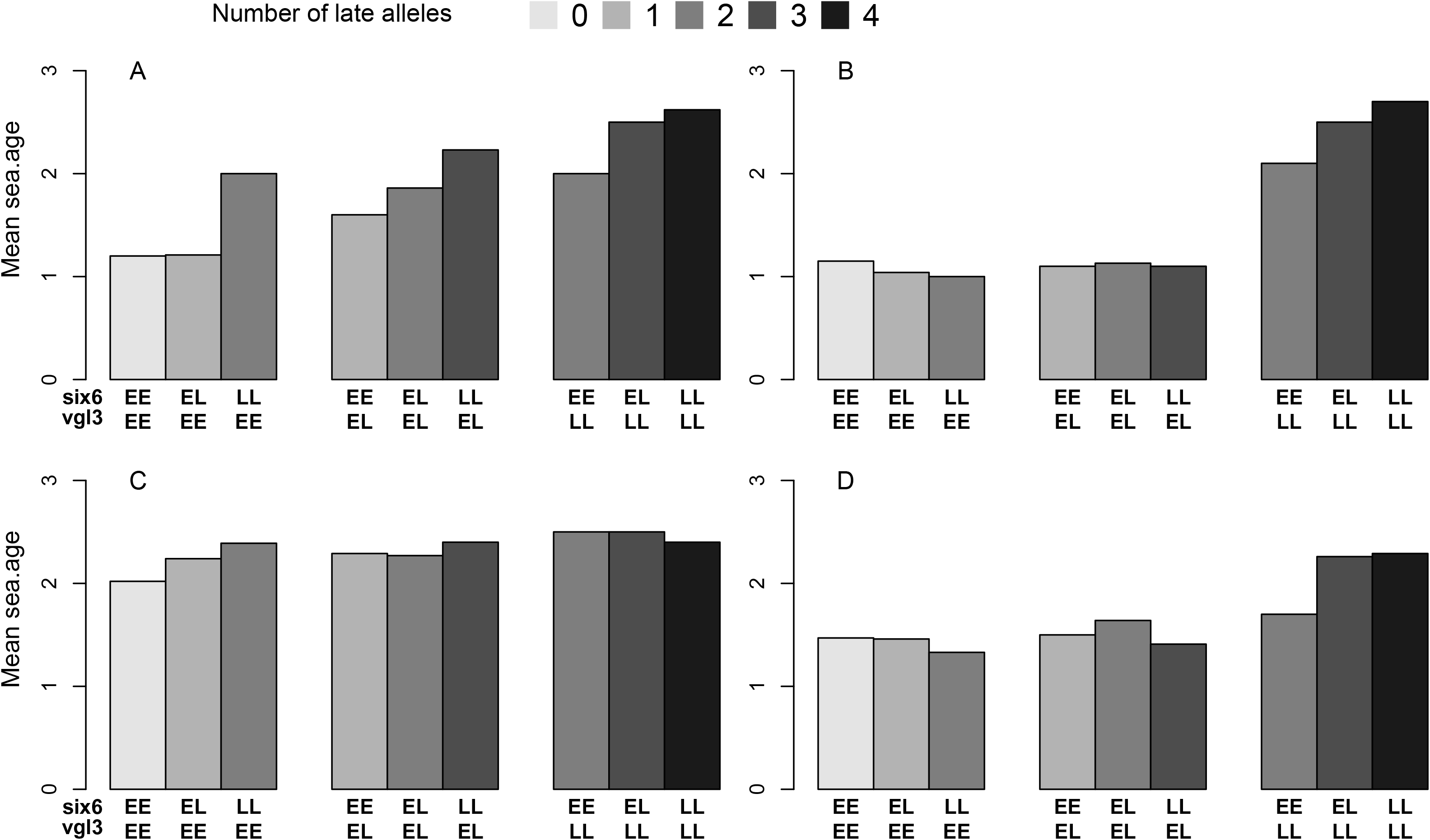
Mean sea age of the samples for each class of genotypes at *vgll3* and *six6*. Females from 1983-84 (**A**), males from 1983-84 (**B**), females from 2013-16 (**C**) and males from 2013-16 (**D**).

On SSA09, four main haplotype sequences linked to the *six6* gene were predominant in both historical and contemporary samples (Table 3). With a respective sea age of 1.3 and 1.4 years for the homozygous fish in the historical samples, both “111” and “211” haplotypes were assigned to the early variant of *six6*, whereas haplotypes “121” and “122”, with a respective mean sea age of 1.9 and 2.1 years, were assigned to the late variant. Following this, we observed an increase in frequency of the *six6* late variant, from 44% in the historical samples to 60% in the contemporary samples (df=1 χ^2^=111, p<2.2.10^−16^). The mean age at maturation in each genotype class (Fig. 2) showed an additive effect of *six6* in the historical female data (Fig. 2.A, Table.S2), and historical male data (Fig. 2.B Table.S2). In contrast, we didn’t observe any association between *six6* and sea age in the contemporary data (Fig. 2.C,D Table.S2).

Cumulatively, *vgll3* and *six6* accounted for 50% of the model deviance in the historical male dataset and 36% in the historical female dataset. In stark contrast, the same loci only accounted for 7% and 3% of the model deviance in the contemporary male and female datasets respectively.

Testing the interaction between these loci revealed significant departure from additivity between *vgll3* and *six6* in the males of the historical samples (Table.S3). This interaction is illustrated in Fig. 2.b where the *six6* genotype is only associated with age at maturation for the samples that are homozygous for the *vgll3-L* allele. No significant departure from additivity was observed in the historical female (Table.S3).

### Genetic admixture

The salmon population inhabiting the river Etne has been subject to introgression from domesticated farmed escapees. We therefore tested whether the observed temporal change in genetic architecture of age at maturation could be linked to admixture with farmed fish. When predicting age at maturation from the joined effect of admixture and genotype, no interaction was detected between individual admixture and *vgll3* genotype (χ^2^=3.6, df=2, p=0.161), nor between individual admixture and *six6* genotype (χ^2^=2.7, df=2, p=0.24). We also checked the allelic frequencies at the *six6* locus in the farmed samples to assess if the increase in *six6-L* allele observed in the contemporary sample could be explained by admixture. With 17% *six6-E* and 68% *six6-L* allele, haplotype frequencies in the farmed samples were more like the contemporary samples than the historical. However, the observed genetic admixture in this population, which is estimated to circa 24%, could only explain a 6% increment in frequency of *six6-*L, which is less than half of the increase observed between historical and contemporary samples.

## Discussion

We document the sudden dissociation between age at maturation in a population of Atlantic salmon and the genotypes of three loci, including the previously identified candidate genes *vgll3* and *six6* (Ayllon *et al*. 2015; Barson *et al*. 2015; Czorlich *et al*. 2018). This dissociation was observed in samples separated by a 30-year time interval and was accompanied by an increase in the frequency of late maturing allele at the *six6* locus, while no change was observed at the *vgll3* locus. The salmon population in the river Etne has been subject to introgression from domesticated salmon escaping from commercial fish farms, with an average 24% of genetic admixture in the contemporary population (Glover *et al*. 2013; Karlsson *et al*. 2016; Besnier *et al*. 2022). However, genetic admixture could not explain alone the change in allelic frequencies observed at *six6*, nor the dissociation between the genes and age at maturation.

The combined effect of *vgll3* and *six6* was not fully additive as we detected a significant epistatic interaction in the historical samples of males. *Six6* was previously identified as a candidate locus associated with age at maturation (Barson *et al*. 2015), but little is known about the genetic effects of this locus. The work presented here is the first to document a sex-specific interaction between *vgll3* and *six6*.

In the river Etne, the frequency of late maturing fish was higher among females than among males. This difference is assumed to be the result of adaptation to sexual conflict where the advantage of maturing late is greater for females than for males (Fleming & Einum 2011). With the *vgll3-E* allele being dominant in males only, *vgll3* is believed to play a key function in the resolution of sexual conflict in the optimal age of maturation in salmon (Ayllon *et al*. 2015; Barson *et al*. 2015). However, with the present results documenting the almost complete dissociation between age at maturation and *vgll3* genotypes in parallel with a trend towards later maturation strategy in both sexes simultaneously, the perennity of local adaptation in the population inhabiting the river Etne can be questioned. Noteworthy, sex-specific strategies of maturation seem to perdure in the population despite *vgll3*’s contribution being almost completely eradicated in the contemporary samples. In fact, the difference in early-maturation frequencies between males and females increases in the contemporary dataset, strongly suggesting that other mechanisms are acting instead of, or in parallel with *vgll3*, to maintain sex specific age at maturation. This hypothesis seems consistent with the description of multiple loci associated with age at maturation (Sinclair-Waters *et al*. 2020).

The absence of association between *vgll3* and age at maturation in multiple North American populations (Boulding *et al*. 2019; Mohamed *et al*. 2019) already suggested that *vgll3* is not the only regulator of sexual conflict for age of maturation in Atlantic salmon. The present study confirms this observation and further shows that the relative influence of *vgll3* is not stable in time. In the case of the population in the river Etne, our data reveals that other unidentified factors also play a major role in maintaining the sex-specific differences in maturation strategies when the link between age at maturation and *vgll3* is disconnected.

The observed trend towards slower marine growth and later maturation in the population inhabiting the river Etne has also been reported in many other salmon populations in the North Atlantic (Quinn *et al*. 2006; Otero *et al*. 2012; Bal *et al*. 2017; Todd *et al*. 2021; Vollset *et al*. 2022). While the precise triggers and mechanisms underpinning to the development of sexual maturation are not yet fully understood in Atlantic salmon (Mobley *et al*. 2021), slow growth and late maturation is consistent with the hypothesis of size or perhaps growth-rate threshold as a determinant for salmon maturation (Rowe *et al*. 1991; Simpson 1992; Taranger *et al*. 2010). Such an energy-budget threshold might also be genetically regulated through the mediation of *vgll3*, which has been shown to be linked with cell fat regulation (Halperin *et al*. 2013) or six6 which has been linked to stomach fullness and prey composition (Aykanat *et al*. 2020). Following this hypothesis, slow growth due to environmental conditions, such as lack of prey would lead to later maturation as a higher proportion of fish would fail to reach the weight (or growth-rate) threshold for maturation after one single year at sea.

Observations suggest important changes in the environmental conditions met by salmon migrating from the river Etne in the period 1980-2010 (Vollset *et al*. 2022). The marine migration pattern for salmon originating from the river Etne is not fully known, however, the first phase is probably a northward migration through the southern Norwegian Sea (Gilbey *et al*. 2021) where changes in the oceanographic conditions have been reported in the last years. For example, water temperature increased by nearly 1°C at 50-200 m depth from early 1980’s until 2021, mainly due to warmer water masses flowing into the southern Norwegian Sea (Skagseth & Mork 2012; ICES 2021a). The early 1980’s are also considered as the end of “The great salinity anomaly”, which was a period with large inflow of cold and fresh Arctic water into the Norwegian Sea. The large proportion of Arctic water entering the Norwegian Sea is correlated to increased productivity (Skagseth *et al*. 2022), and to improved feeding conditions for post-smolts in the region (Utne *et al*. 2022). Therefore, salmon returning in 1983 and 1984 had probably been feeding in a very productive sea during the initial post-smolt phase, whereas salmon returning to rivers in 2013-2016 had been feeding in a warm and saline Norwegian Sea. During this later period, observed stomach fullness and condition factor for post-smolt sampled in the Norwegian Sea were low (Utne *et al*. 2021a). In addition, the potential interspecific competition with other pelagic fish for prey (Utne *et al*. 2021b) was low in the early 1980’s as the total biomass of pelagic fish feeding in the Norwegian Sea in 1982-1983 was around 1/3 of the total biomass in the period 2013-2016 (ICES 2008, 2021b) when Norwegian Spring-spawning herring had not yet recovered from the collapse in the late 1960ies, and NEA-mackerel and blue whiting stock biomasses were at low levels (ICES 2008). Many observations seem to confirm that environmental conditions were less favorable for salmon growth in the last decade than in the early 1980’s. It is thus conceivable that the changes observed in the river Etne are caused by environmental perturbations occurring over a large region and affecting salmon populations as well as other organisms.

Under commercial aquaculture conditions, farmed Atlantic salmon typically display early sexual maturation (Taranger *et al*. 2010), and importantly, the relative influence of *vgll3* on age at maturation appears to be largely bypassed (Ayllon *et al*. 2019). Yet again, this response is sex-specific as a study conducted under aquaculture conditions (Ayllon *et al*. 2019) reported that *vgll3* did not show any correlation with age at maturation in females while displaying only weak association in males (Ayllon *et al*. 2019). The commercial farm strain used in the aforementioned study, known as Mowi, stemmed from wild Norwegian salmon populations in which *vgll3* was identified as a candidate gene for influencing age at maturation (Ayllon *et al*. 2015; Barson *et al*. 2015). The authors concluded that high calorie feed intake combined with artificial light and temperature regimes as well as potential genetic or epigenetic components, may alter the impact of *vgll3* on age at maturation (Ayllon *et al*. 2019). Anecdotally, it is also worth noting that despite multiple generations of directional selection against early maturing fish in the domesticated farmed salmon, the genetic variability of *vgll3* remains high in the Mowi strain (Ayllon *et al*. 2019). It is thus possible that a rapid change in environmental conditions, such as feed intake and therefore growth-rate, may bypass the effect of *vgll3* without letting selection (artificial selection in this case) alter the allelic frequencies of the gene.

We thus can hypothesize a two-threshold model where the effect of *vgll3* on age at maturation is bypassed when abundance of feed resources is very high, as in farming conditions, or when feed resources are very low and fish need to spend more time at sea to acquire the necessary energy-reserves for maturation and reproduction. When resource availability falls between both thresholds, *vgll3* may play a more important role in determining age at maturation. Inversely if feed resources exceed the high threshold, or do not reach the low one, *vgll3* is bypassed by environmental conditions, and therefore age at maturation is determined by a combination of other genetic and environmental factors.

In the face of changing environmental conditions, one may expect the allelic frequencies at the associated loci to change in response to the newly induced selection pressure (Czorlich *et al*. 2018; Jensen *et al*. 2022). This is the case for *six6* where the temporal trend towards later maturation is accompanied by an increase of the *six6-*L allele frequency. This shift in allelic frequencies is likely due to positive selection on the *six6-L* allele, alone, or in conjunction with admixture from domesticated salmon where *six6-*L was found in higher frequency. In contrast, the *vgll3* allelic frequencies remained stable despite major changes in age at maturation. This result seems to indicate a change in environmental conditions, creating a situation where the influence of *vgll3* was effectively bypassed before natural selection had time to operate, whereas selection had the time to modify the allelic frequencies at the *six6* locus before the influence of this gene was also bypassed. This hypothesis is strongly supported by the observations of sudden change in age at maturation in many Atlantic salmon populations inhabiting Northeast Atlantic rivers (Vollset *et al*. 2022), including the population in the river Etne (Harvey *et al*. 2022), where a highly distinct and sudden drop in early marine growth was reported in 2005 and subsequently, referred to as an ecological regime shift in the northeast Atlantic ocean (Vollset *et al*. 2022).

As this, and other salmon population in the Northeast Atlantic were subjected to the same ecological regime shift, leading to consistently reduced marine growth rates and increased age at maturation, we conclude that growth-driven plasticity has almost completely bypassed the combined influence of *vgll3* and *six6* within on age at maturation, and furthermore, on resolving the sexual conflict for this trait in Atlantic salmon. Together with the interaction between *vgll3* and *six6* described in the historical data, the dissociation between genes and age at maturation represents an original finding that changes our understanding of the genetic architecture of age at maturation in Atlantic salmon.

## Acknowledgments

This work was financed by the Norwegian Ministry of Trade, Industry and Fisheries. This authority paid no part in the study design nor interpretation of results. We would like to acknowledge the research institute Nina, and anglers, for donating some of the farmed salmon samples used in this study. We would like to acknowledge the river Etne owners association for continued collaboration and allowing capture and handling of fish and IMR staff involved in monitoring the population in the trap. We would like to thank María Quintela for insightful comments on the earlier drafts of the manuscript, and Emily K. Glover for drawing the salmons used in Figure 1.

## Data availability

Data for this study are available from the IMR.brage.unit.no public repository. Link to data: https://imr.brage.unit.no/

