## supplementary tables 1 to 3 for "Overruled by nature: A plastic response to an ecological regime shift disconnects a gene and its trait"

| Coefficients: | estimate | Std. Error | z value | Pr(>\|z\|) |
| --- | --- | --- | --- | --- |
| Females 1983-84 | |  |  |  |
| Intercept | 6.75 | 0.86 | 7.81 | <0.001 |
| Year(1984) | -1.81 | 0.34 | -5.27 | <0.001 |
| Vgll3 (a) | -2.26 | 0.33 | 6.78 | <0.001 |
| Vgll3 (d) | -0.02 | 0.39 | -0.07 | 0.941 |
| Males 1983-84 | |  |  |  |
| Intercept | 9.00 | 0.94 | 9.5 | <0.001 |
| Year(1984) | -1.62 | 0.39 | -4.13 | <0.001 |
| Vgll3 (a) | -2.73 | 0.30 | -4.13 | <0.001 |
| Vgll3 (d) | 1.91 | 0.36 | 5.26 | <0.001 |
| Females 2013-16 | |  |  |  |
| Intercept | 17.07 | 2252.43 | 0.01 | 0.99 |
| Year(2014) | -0.25 | 0.48 | -0.51 | 0.60 |
| Year(2015) | -0.26 | 0.57 | -0.46 | 0.64 |
| Year(2016) | -18.56 | 1684.11 | -0.01 | 0.99 |
| Vgll3 (a) | -9.22 | 1126.21 | -0.01 | 0.99 |
| Vgll3 (d) | 8.46 | 1126.21 | 0.01 | 0.99 |
| Males 2013-16 | |  |  |  |
| Intercept | 2.54 | 0.62 | 4.08 | <0.001 |
| Year(2014) | -0.02 | 0.31 | -0.08 | 0.93 |
| Year(2015) | 1.59 | 0.33 | 4.7 | <0.001 |
| Year(2016) | -0.90 | 0.37 | -2.43 | 0.015 |
| Vgll3 (a) | -1.06 | 0.23 | -4.59 | <0.001 |
| Vgll3 (d) | 0.65 | 0.28 | 2.33 | 0.020 |

Table S1. **Summaries of binomial GLMs of sea age**. Models test the effect of capture year, vgll3 additive (a) and dominance (d). Models were fitted separately for males and females, historical and contemporary samples.

Table S2 **Summaries of binomial GLMs of sea age**. Models test the effect of capture year and six6 additive (a). Models were fitted separately for males and females, historical and contemporary samples

| Coefficients: | estimate | Std. Error | z value | Pr(>\|z\|) |
| --- | --- | --- | --- | --- |
| Females 1983-84 | |  |  |  |
| Intercept | -2.92 | 0.66 | -4.37 | <0.001 |
| Year(1984) | -1.14 | 0.25 | -4.43 | <0.001 |
| Six6 (a) | 1.06 | 0.21 | 5.07 | <0.001 |
| Males 1983-84 | |  |  |  |
| Intercept | -0.73 | 0.61 | -1.20 | 0.22 |
| Year(1984) | -1.09 | 0.24 | -4.43 | <0.001 |
| Six6 (a) | 0.77 | 0.20 | 3.79 | <0.001 |
| Females 2013-16 | |  |  |  |
| Intercept | -2.92 | 0.97 | -3.01 | 0.002 |
| Year(2014) | -0.25 | 0.48 | -0.51 | 0.60 |
| Year(2015) | -0.29 | 0.56 | -0.52 | 0.60 |
| Year(2016) | -17.68 | 1064.83 | -0.01 | 0.99 |
| Six6 (a) | 0.37 | 0.31 | 1.15 | 0.24 |
| Males 2013-16 | |  |  |  |
| Intercept | 0.01 | 0.52 | 0.01 | 0.99 |
| Year(2014) | -0.07 | 0.30 | -0.02 | 0.98 |
| Year(2015) | 1.71 | 0.32 | 5.2 | <0.001 |
| Year(2016) | -0.86 | 0.35 | -2.41 | 0.015 |
| Six6 (a) | -0.03 | 0.17 | -0.20 | 0.84 |

Table S3. Summaries of binomial GLMs of sea age. Models test the effect of capture year and vgll3 (a), six6 additive (a), and the interaction vgll3*six6. Models were fitted separately for males and females, historical and contemporary samples

| Coefficients: | estimate | Std. Error | z value | Pr(>\|z\|) |
| --- | --- | --- | --- | --- |
| Females 1983-84 | |  |  |  |
| Intercept | 3.03 | 3.48 | 0.87 | 0.38 |
| Year(1984) | -1.81 | 0.34 | -5.21 | <0.001 |
| Vgll3 (a) | -1.68 | 1.30 | -1.30 | 0.19 |
| Six6 (a) | 1.11 | 1.08 | 1.02 | 0.31 |
| Vgll3 (a) * Six6 (a) | -0.15 | 0.41 | -0.37 | 0.71 |
| Males 1983-84 | |  |  |  |
| Intercept | 23.68 | 4.66 | 5.07 | <0.001 |
| Year(1984) | -1.50 | 0.34 | -4.42 | <0.001 |
| Vgll3 (a) | 6.95 | 1.44 | -4.81 | <0.001 |
| Six6 (a) | -4.24 | 1.33 | -3.16 | 0.001 |
| Vgll3 (a) * Six6 (a) | 1.33 | 0.42 | 3.12 | 0.001 |
| Females 2013-16 | |  |  |  |
| Intercept | -2.24 | 4.22 | -0.52 | 0.59 |
| Year(2014) | -0.23 | 0.49 | -0.47 | 0.63 |
| Year(2015) | -0.22 | 0.57 | -0.39 | 0.69 |
| Year(2016) | -17.57 | 1028.71 | -0.01 | 0.89 |
| Vgll3 (a) | -0.23 | 1.65 | -0.14 | 0.88 |
| Six6 (a) | 1.07 | 1.45 | 0.74 | 0.45 |
| Vgll3 (a) * Six6 (a) | -0.29 | 0.57 | 0.50 | 0.61 |
| Males 2013-16 | |  |  |  |
| Intercept | 5.55 | 2.41 | 2.30 | 0.02 |
| Year(2014) | -0.05 | 0.32 | -0.18 | 0.85 |
| Year(2015) | 1.64 | 0.33 | 4.9 | <0.001 |
| Year(2016) | -0.95 | 0.83 | -2.35 | 0.01 |
| Vgll3 (a) | -1.95 | 0.83 | -2.35 | 0.02 |
| Six6 (a) | -1.15 | 0.82 | -1.40 | 0.16 |
| Vgll3 (a) * Six6 (a) | 0.39 | 0.28 | 1.38 | 0.16 |
